# Generation of transducible version of a recombinant human HAND2 transcription factor from *Escherichia coli*

**DOI:** 10.1101/2021.09.04.458986

**Authors:** Krishna Kumar Haridhasapavalan, Pradeep Kumar Sundaravadivelu, Anshuman Mohapatra, Neha Joshi, Nayan Jyoti Das, Rajkumar P Thummer

## Abstract

Transcription factor HAND2 has a significant role in vascularization, angiogenesis, and cardiac neural crest development. Also, it is one of the key cardiac factors crucial for the enhanced derivation of functional and mature myocytes from non-myocyte cells. Here, we report the generation of the recombinant human HAND2 fusion protein from the heterologous system. First, we cloned the full-length human *HAND2* gene (only protein-coding sequence) after codon optimization along with the fusion tags (for cell penetration, nuclear translocation, and affinity purification) into the expression vector. We then transformed and expressed it in *Escherichia coli* (*E. coli*) strain, BL21(DE3). Next, the effect (in terms of expression) of tagging of fusion tags with this recombinant protein at two different terminals was also investigated. Notably, using affinity chromatography, we established the one-step homogeneous purification of human recombinant HAND2 protein; and through circular dichroism spectroscopy, we established that this purified protein had retained its secondary structure. Furthermore, we show that this purified human protein could transduce the human cells and translocate to its nucleus. Prospectively, the purified recombinant HAND2 protein can potentially be a safe and effective molecular tool in the direct cardiac reprogramming process and other biological applications.

## Introduction

The role of recombinant proteins in therapeutics has been indispensable for the past four decades and will continue to be so. Production of recombinant proteins using different host organisms such as algae [1], bacteria [2], yeast [3], insects [4], and mammalian cells [5] has so far proven to be a complex but effective process [6]. Therefore, an immense number of recombinant proteins have been expressed, purified, and used for a wide range of biotechnological applications [7]. The most commonly used expression host system for recombinant protein production is the bacterial system, especially *Escherichia coli (E. coli)* strains due to easy handling and maintenance, well-studied genetics, well-understood cell machinery, high protein yield, and so forth [8–10]. Once expressed, these proteins are purified using a wide range of purification tags. The most widely used tag for purification is the poly-histidine tag since it is inexpensive and does not alter the characteristics of the proteins [11, 12]. The introduction of recombinant proteins into mammalian cells has been proven to be an effective alternative since it does not integrate and alter the genome, and also manipulation of cell fate can be done in a time and dosage-dependent manner [13–16]. Thus, recombinant proteins contribute to a major and vital part in therapeutics and in safer and non-integrative cell reprogramming processes. However, various challenges are associated with the successful production of these therapeutic recombinant proteins, such as codon usage bias, gene toxicity, mRNA instability, poor protein expression, proteolytic cleavage by the host cell, purity, poor solubility and stability (*in vitro*), and protein misfolding [13, 17]. In this study, by circumventing these limitations, we aimed to generate a cell- and nuclear-permeant human Heart- and neural crest derivatives-expressed protein 2 (HAND2) protein that can prospectively be used for various biological applications.

HAND2 is a member of the basic helix-loop-helix family of transcription factors that have a consensus DNA binding sequence 5’-CANNTG-3’ [18]. The transcript of the HAND2 gene is 2780 bp long, containing two exons. The coding sequence is 654 bp in length which translates to the protein of 218 amino acids. HAND2 is highly expressed in maternal decidua [19] and adult heart, liver, and testes [19]. The knockout of the *HAND2* gene in mice leads to defects in ventricle formation, eventually leading to embryonic lethality [18, 20]. A recent study reported HAND2 regulated genes involved in the atrioventricular canal and cardiac valve development [21]. Besides, HAND2 governs the development of epicardium, vascularization and angiogenesis, second heart field development and survival, and cardiac neural crest development [22]. All these studies indicate the role of HAND2 in cardiac development, both in mesoderm-derived and neural crest-derived structures.

Apart from these, HAND2 plays a critical role in the development of other tissues during mouse embryogenesis. It is known to have a significant role in the development of branchial arch and limb bud [20, 23]. HAND2 regulates the anterior-posterior polarity of limb bud through a chain of downstream transcriptional regulators [23]. It also aids in craniofacial development [20], and is essential for developing the sympathetic nervous system in humans, especially noradrenergic neurons [24].

In addition, HAND2 is crucial for the formation and maturation of cardiomyocytes [18]. Therefore, it is regarded as one of the most crucial cardiac reprogramming factors to derive functional cardiomyocytes. Several studies have used the genetic form of HAND2 in their reprogramming cocktail, and its inclusion resulted in higher efficiency than the original GATA4, MEF2C, and TBX5 combination [25–27]. Recently, the role of HAND2 in altering the chromatin accessibility and gene expression in fibroblasts to convert them to cardiac pacemaker-like cells has also been reported [28].

Recently, we have demonstrated the heterologous expression and purification of human cardiac reprogramming factors, namely GATA4 [29], ETS2 [30], and MESP1 [31], in recombinant forms. Here, we demonstrated the soluble expression and purification of recombinant human HAND2 protein from *E. coli* under native conditions having efficient cell permeability and nuclear translocation ability. This is the first study to identify the optimal induction parameters for the soluble expression and purify (native conditions) this transcription factor, HAND2, from *E. coli*.

## Materials and methods

### Construction of *HAND2* fusion genetic constructs

The protein-coding sequence of the *HAND2* gene was retrieved, codon-optimized, evaluated, and cloned in a protein expression vector as depicted in Figure S1. The resulting plasmids were verified by DNA sequencing and also by restriction digestion analysis using different combinations of restriction enzymes. The *in silico* online tools used for codon optimization and its evaluation are listed in Table S1. The primers used for sequencing are listed in Table S2.

### Identification of ideal expression parameters

*E. coli* BL21(DE3) cells were used for all the expression analysis experiments and purification. The cells were transformed with appropriate recombinant plasmids harboring the *HAND2* fusion gene and cultured as described recently [31]. Subsequently, the culture (20 mL) was induced with a required concentration of Isopropyl β-D-1-thiogalactopyranoside (IPTG) (HiMedia) and incubated for a respective time at a respective temperature depending on experimental requirements. The various values of the parameters such as IPTG concentration, pre-induction cell density, postinduction incubation time, and induction temperature screened for analysis are listed in Table S3. Uninduced cultures (20 mL) were also incubated as comparable experimental conditions. After the incubation, the cells were harvested and lysed in prechilled lysis buffer (1.2 mL) (Table S4) by ultrasonication on ice. The obtained total cell lysate was clarified to separate the insoluble pellet and the soluble supernatant fractions. All the protein samples were analyzed using sodium dodecyl sulfate-polyacrylamide gel electrophoresis (SDS-PAGE) and immunoblotting.

### Immobilized Metal Ion Affinity Chromatography

From the soluble fraction, the human recombinant HAND2 fusion protein was purified using Immobilized Metal Ion Affinity Chromatography under native conditions. Briefly, HTN-HAND2 expression was induced in large culture volumes (600 mL) with optimized expression parameters. Subsequently, cells were harvested, resuspended in lysis buffer (20 mL), and further lysed using ultrasonication on ice. The total crude cell lysate was clarified by centrifugation to obtain a soluble supernatant fraction. The soluble fraction was incubated with nickel-nitrilotriacetic acid (Ni-NTA) resin at 4 °C for 8 to 14 h with continuous shaking. After incubation, the soluble fraction with Ni-NTA was loaded onto the purification column (Bio-Rad), equilibrated with lysis buffer, and the flow-through was drained out. Consequently, the column was washed with wash buffers (100 mL) sequentially. Once the wash buffers were fully drained out, elution buffer was applied to elute out the desired HAND2 recombinant protein. The PD10 columns (GE Healthcare) were used to desalt the purified protein according to the manufacturer’s instructions against glycerol buffer and stored at -80 °C until further use. The purity of the recombinant HAND2 protein was established by SDS-PAGE and identity was confirmed with immunoblotting. All the buffers used in this purification, along with their compositions, are listed in Table S4. The pH of the buffers was adjusted to 8.0 at room temperature and prechilled on ice before use.

### SDS-PAGE and immunoblotting

The protein concentration of sonicated and purified samples was estimated using Bradford assay [32]. The SDS-PAGE and immunoblotting were performed as described in our recent studies [29, 30]. The antibodies used and their respective concentrations are listed in Table S5.

### Far-ultraviolet circular dichroism spectroscopy

Far-ultraviolet (UV) circular dichroism (CD) spectroscopy was carried out as described in our recent studies [29, 30]. From the CD spectrum obtained, the secondary structure content of the purified HAND2 protein was estimated using the online tool, Beta Structure Selection (BeStSel; http://bestsel.elte.hu/index.php).

### MALDI-TOF

The purified protein was desalted in deionized Milli-Q water using a PD-10 column from GE healthcare and concentrated fivefold using a Thermo scientific three kDa MWCO protein concentrator. The protein was diluted two times in α-Cyano-4-hydroxycinnamic acid matrix solution. The molecular weight was accurately determined using the Bruker Autoflex speed MALDI-TOF mass spectrometer.

### Cell culture

Human foreskin fibroblasts (HFF), BJ, (ATCC® CRL-2522™) were cultured (in gelatin-coated dishes) in fibroblasts growth medium (FGM) containing 10% fetal bovine serum (FBS; Invitrogen), 1X non-essential amino acids (NEAA; Invitrogen) and 1% penicillin/streptomycin (P/S; Invitrogen) in high-glucose DMEM (Invitrogen) at standard cell culture conditions (37 °C with 5% CO_2_ in a humidified atmosphere). This cell line was procured from American Type Culture Collection, USA. HeLa cells were cultured in HeLa growth medium (HGM) containing 10% FBS, and 1% P/S in high-glucose DMEM at standard cell culture conditions. This cell line was procured from National Centre for Cell Science, Pune, India. Both these cells were passaged with trypsin-EDTA (Invitrogen) at ∼80-90% confluency in the ratio 1:4.

### HAND2 fusion protein transduction

HFFs and Hela cells were adjusted to 5 x 10^4^ and 1 x 10^5^ cells/well, respectively, with growth medium and seeded in 24-well culture plates. Cells were grown 12-24 h at 37 °C with 5% CO_2_ under humidified culture conditions and then treated with protein transduction medium (DMEM, 2% FBS, 1X NEAA, 1% P/S, and optimal concentrations of purified recombinant HTN-HAND2 protein or glycerol buffer as a control) and re-incubated for 4-6 h. Post-incubation, cells were washed with phosphate buffer saline and used for further analysis.

### Immunocytochemistry and microscopy

Immunocytochemistry and microscopy were performed as described recently [29, 33]. For immunocytochemistry, the antibodies and their respective concentrations used are listed in Table S5. For microscopy, the images were acquired using an inverted fluorescence microscope (IX83, Olympus, Japan) equipped with a DP80 CCD camera. Samples were illuminated using a pE-300 white CoolLED light source. Cells were then counterstained with DAPI (Sigma), and image stacks were acquired using the 20x/0.45NA objective at 2 μm intervals. Images were analyzed by CellSens dimension (Olympus) and Image J software.

### Nucleotide sequence

The codon-optimized (for expression in *E. coli* strain BL21(DE3)) nucleotide sequence of the *HAND2* gene can be retrieved from the NCBI GenBank database (accession code MW570765).

## Results

### Codon optimization and cloning of human *HAND2* gene sequence

We first codon-optimized the non-optimized *HAND2* protein-coding sequence for expression in *E. coli*. The alignment of non- and codon-optimized codons and their respective amino acids is depicted in Fig. 1. Before and after codon optimization, the nucleotide sequence was analyzed using two different online tools for comparison. When analyzed using the Genscript Rare Codon Analysis (GRCA), the non-optimized sequence had approximately 7% of codons that might have hindered the expression, which were then replaced after codon optimization with the codons that are abundantly used in *E. coli* (Figure S2; Table 1). A similar analysis was performed using Graphical Codon Usage Analyzer (GCUA), and it was found that seven codons were present in the non-optimized sequence (Figure S3; *in magenta*) that might have hindered the expression, which were then replaced with the codons that are abundantly used in *E. coli* (Figure S3; *right*). As an example, the CTC rare codon present in the first 50 codons of the sequence [Fig. S3; top (*in magenta*)] was substituted with CTG codon [Fig. S3; top (*in black*)]. Moreover, codon adaptation index value improved from 0.69 for non-optimized to 0.89 for codon-optimized sequence (Table 1). Ideally, the codon adaptation index value should lie between 0.8 and 1.0 (Table 1), which indicates that the optimized sequence of *HAND2* is ideal for expression in *E. coli*.

**Fig. 1.**
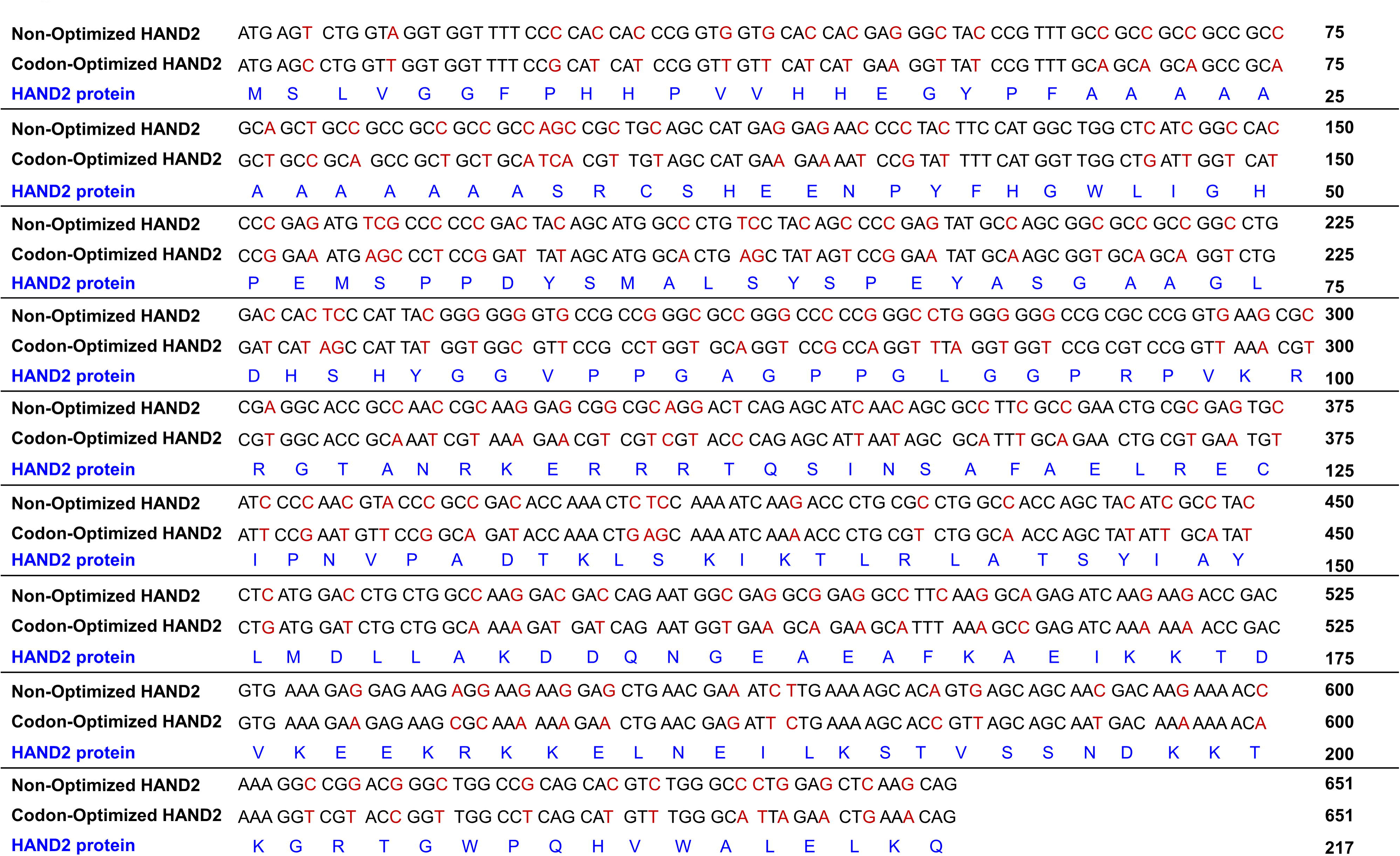
Comparison of *HAND2* gene sequences before and after codon optimization. The codons before and after optimization are aligned with the respective amino acids. The change in nucleotides is represented in red.

**Table 1.** Parametric GRCA tool of non-optimized as well as codon-optimized coding sequence of human *HAND2* gene

| Parameters | Non-optimized | Codon-optimized | Ideal range |
| --- | --- | --- | --- |
| Codon Adaptation Index | 0.69 | 0.89 | 0.8-1.0 |
| GC Content | 67% | 49% | 30-70% |
| Codon Frequency Distribution | 7% | 0% | <30% |

Next, we tagged this codon-optimized *HAND2* gene sequence with a set of fusion tags (His, TAT, and NLS) at either end to generate *HAND2* fusion gene inserts [Fig. 2a and 2b (*top*)]. Various studies have reported that the expression, solubility, and stability of human proteins heterologously expressed in *E. coli* is influenced by the position of the fusion tags [29-31, 33-36]. These fusion gene inserts (HTN-*HAND2* and *HAND2*-NTH) were artificially synthesized and subcloned into the pET28a(+) expression vector containing a tightly regulated T7 promoter (inducible). Restriction digestion with selected restriction enzymes and subsequent fragments on agarose gel verified the successful cloning of the *HAND2* fusion gene inserts into the vector [Fig. 2a and 2b (*bottom*); Table S6]. Further, we confirmed the successful cloning of the *HAND2* fusion genes in the proper orientation and also the fidelity of its nucleotide sequences by DNA sequencing (data not shown). Thus we developed the recombinant pET28a(+) plasmids harboring *HAND2* fusion genes [pET28a(+)-HTN-*HAND2* and pET28a(+)-*HAND2*-NTH constructs], which can be used to produce HAND2 fusion proteins (HTN-HAND2 and HAND2-NTH) with subcellular and subnuclear localization ability.

**Fig. 2.**
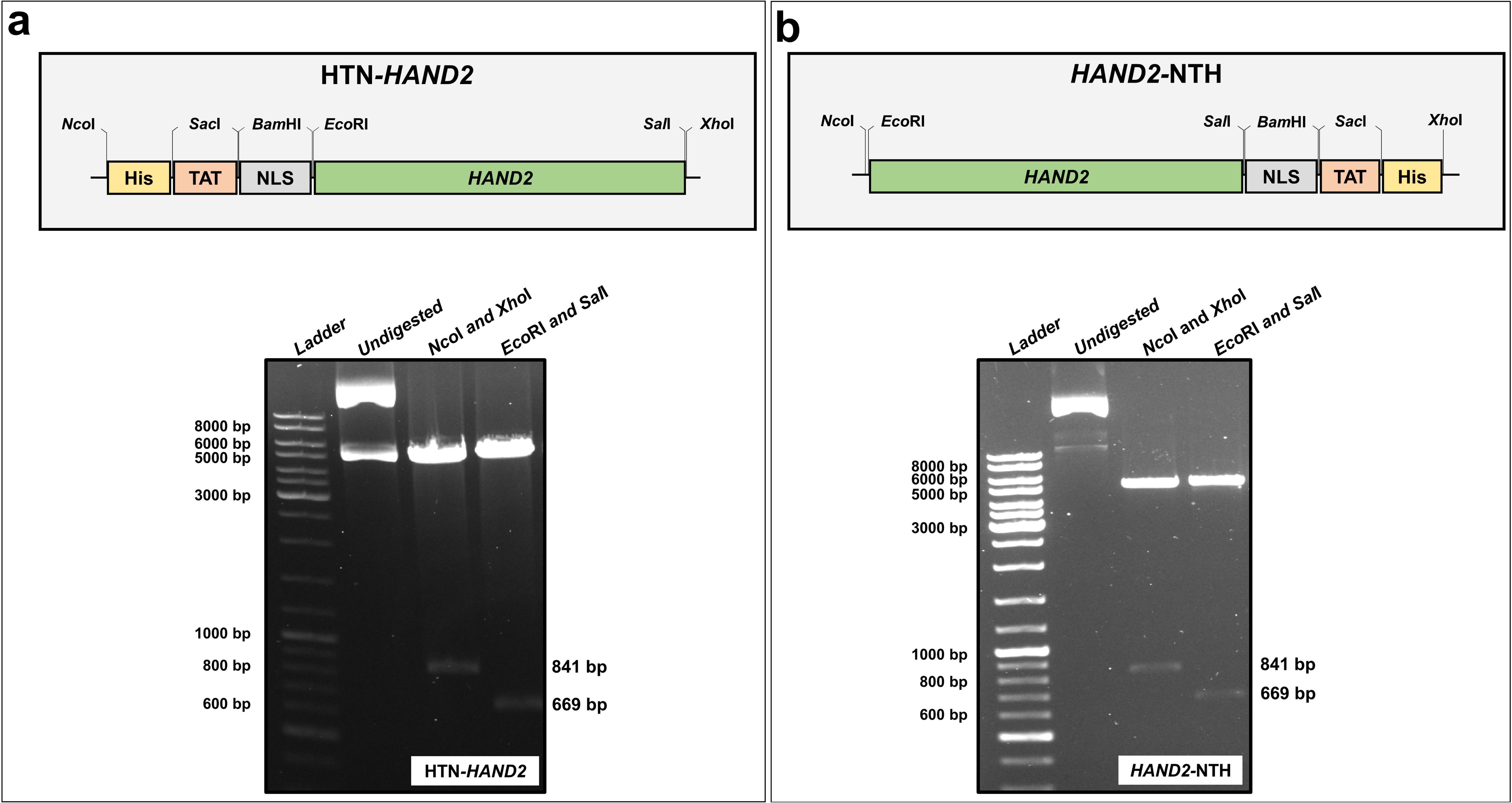
Schematic illustration of the tagging pattern in human *HAND2* gene and verification of cloning. **a** An Illustration of HTN-*HAND2* (*top*) gene insert (not drawn to scale) and verification of the cloned plasmid pET28a(+)-HTN-*HAND2* (*bottom*) using restriction digestion analysis. **b** Illustration of *HAND2*-NTH (*top*) gene insert (not drawn to scale) and verification of the cloned plasmid pET28(+)-*HAND2*-NTH (*bottom*) using restriction digestion analysis. His (H): polyhistidine (8X); TAT (T): Transactivator of Transcription; NLS (N): Nuclear Localization Sequence/Signal.

### Screening various parameters for obtaining soluble expression of the recombinant HAND2 fusion protein

We next sought to determine optimal parameters for obtaining maximal soluble expression. Numerous studies have identified the optimal (ideal) conditions for achieving the maximal soluble expression of bioactive recombinant proteins [17, 37–43]. In the process, we screened various parameters, namely pre-induction cell density, inducer concentration, and postinduction incubation time. The different values that were screened for each parameter are listed in Table S3. Firstly, from the screening experiments, optimal pre-induction cell density, inducer concentration, and postinduction incubation time was found to be ∼0.5 OD_600_, 0.05 mM, and 2 h, respectively. Subsequently, we examined the soluble expression of both the fusion proteins (HTN-HAND2 and HAND2-NTH) induced at 37 °C, using SDS-PAGE and immunoblotting. These results (Fig. 3A and 3B) revealed that in both the cases (HTN-HAND2 and HAND2-NTH), nearly half of the overall expressed HAND2 protein molecules are found in pellet/insoluble cell fractions and rest in the supernatant/soluble cell fractions. Earlier reports recommended that reduction in induction temperature enhances the protein solubility [37, 39, 42]. Therefore, we further investigated whether the reduction in temperature enhances the solubility of the HAND2 fusion protein. This is crucial to avoid purification from inclusion bodies, which contains partially folded or unfolded protein of interest and requires strong detergents to solubilize the inclusion bodies and extract the protein of interest and then refold to its native state [17, 44]. In both cases, induction at 18 °C considerably improved the solubility of the recombinant HAND2 fusion protein (Fig. 3a and 3b). However, a decline in the overall expression only in HAND2-NTH fusion protein was observed when induced at 18 °C compared to 37 °C. The immunoblotting analysis further represents the truncated protein fragments of HAND2-NTH protein in both the induction temperatures, unlike HTN-HAND2 [Fig. 3a and 3b (*bottom*)]. These truncated fragments could be due to various possible reasons such as: (i) intragenic sequences that mimic *E. coli* ribosomal entry sites present within the protein-coding sequence [45], (ii) proteolysis of some protein molecules at specific sensitive sites during expression [46], (iii) protein cleavage at Asp-Pro bonds because of overheating of protein samples [47]. Among the three reasons, the last reason is least likely in this study as the protein samples of both HTN-HAND2 and HAND2-NTH were treated similarly, and no such truncations were observed in the case of HTN-HAND2. Importantly, these truncations might compromise the full-length recombinant HAND2 fusion protein quality and purity at the final stages. Moreover, this observation signified the importance of induction temperature and the terminal at which the fusion tags were coupled in the production of quality recombinant proteins. An earlier study also reported similar observations with different transcription factors [35]. Thus, based on this analysis, HAND2 protein with fusion tags at the N-terminal end of HAND2 (HTN-HAND2) induced at 37 or 18 °C, were selected for further experiments.

**Fig. 3.**
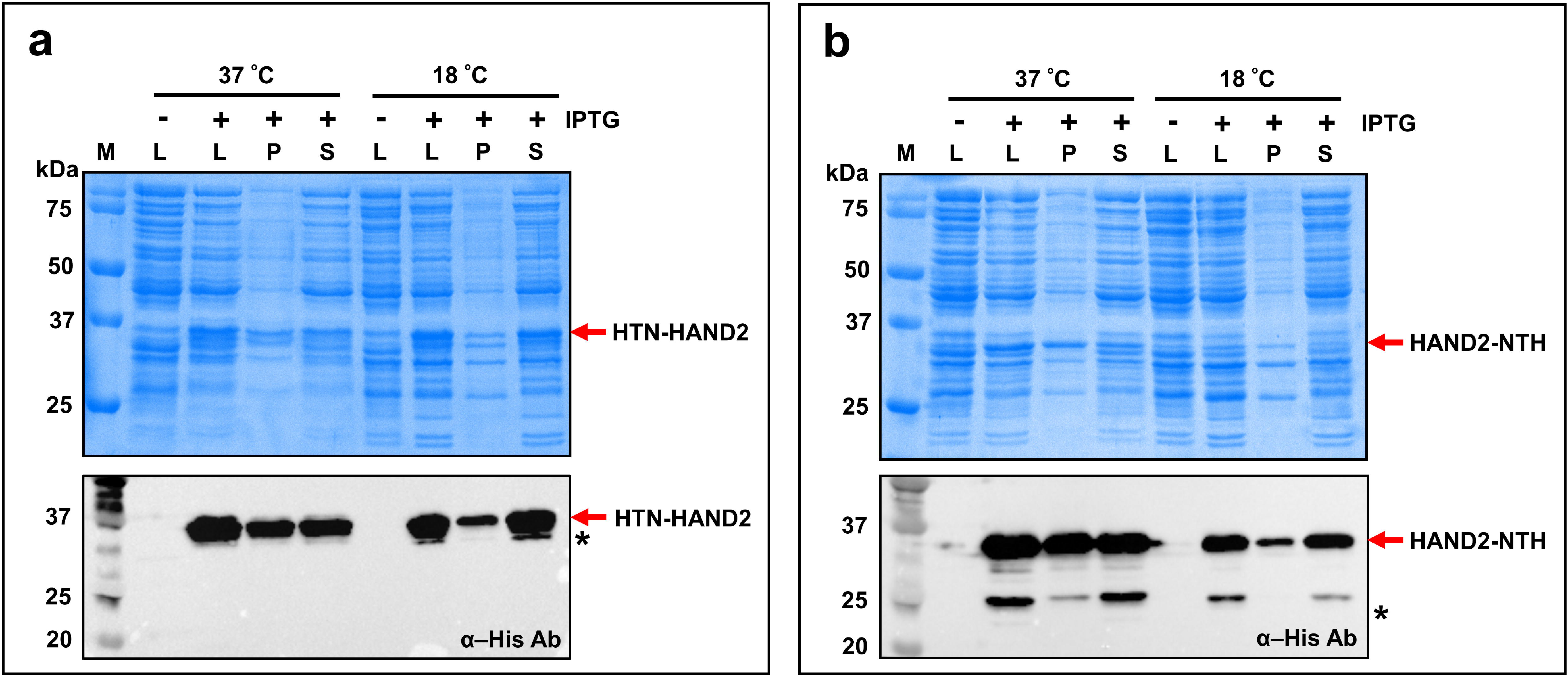
Analyzing the effect of induction temperature and tagging pattern on the expression of human HAND2 protein. *E. coli* BL21(DE3) cells were transformed with pET28a(+)-HTN-*HAND2* **a** and pET28a(+)-*HAND2*-NTH **b** and then induced and incubated either at 37 °C for 2 hours or 18 °C for 24 hours. The cell lysate, soluble, and insoluble cell fractions obtained by lysing the cells were analyzed by SDS-PAGE (*top*) and immunoblotting (*bottom*) with normalized loading. (n=2). *Truncations of the HAND2 fusion protein. M: Protein marker (kDa); L: Total cell lysate; P: Pellet/insoluble cell fraction; S: Supernatant/soluble cell fraction; Ab: Antibody.

### One-step purification of the recombinant human HAND2 fusion protein

To purify recombinant HTN-HAND2 protein from soluble cell fraction, IMAC (in this study, Ni^2+^-NTA resin) under native conditions was performed to retain the native-like secondary structure conformation of this fusion protein. Markedly, proteins purified under native (soluble cell fraction) conditions often produced bioactive molecules with native-like folding conformations [48]. At first, the effect of imidazole concentration on the elution of HAND2 fusion protein, induced at 37 °C, was investigated. As shown in Fig. 4a, this recombinant fusion protein started eluting with 200 mM of imidazole, and the maximum amount was eluted with 250-350 mM of imidazole. Importantly, no bacterial proteins were observed in any of the elution. The imidazole gradient elution profile of affinity-purified HAND2 protein, induced at 37 °C, is shown in Fig. 4b. Thus, for the one-step purification of the recombinant HAND2 fusion protein, induced at two different temperatures, we used a maximum of 150 mM of imidazole during washing and 300 mM of imidazole for eluting the HAND2 protein.

**Fig. 4.**
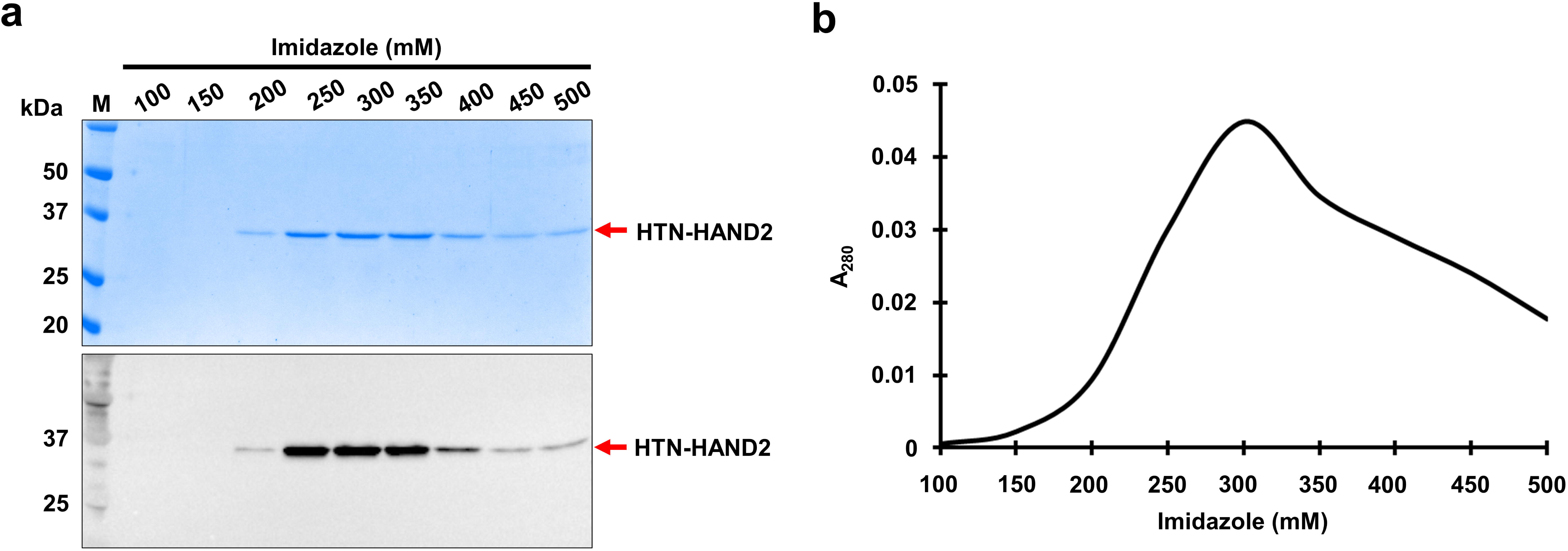
Effect of imidazole concentration on the amount of HAND2 protein eluted. **a** The expressed HTN-HAND2 protein was purified using affinity chromatography. During purification, the imidazole concentration in the elution buffer was varied to see its effect on the amount of protein eluted. As mentioned in the figure, different concentrations of imidazole were used in the elution buffer, and the protein was eluted. The eluted proteins were analyzed using SDS-PAGE (*top*) and immunoblotting (*bottom*) using an α-his antibody. **b** The absorbance of the protein eluted using different concentrations of imidazole in elution buffer were measured at 280 nm, and a graph was plotted with imidazole concentration on the x-axis against absorbance at 280 nm on the y-axis.

With the identified expression conditions, we expressed and purified the recombinant HTN-HAND2 protein under native conditions, as shown in Fig. 5a. From the SDS-PAGE and immunoblotting analysis, it is clear that the recombinant HTN-HAND2 protein, induced at two different temperatures, 37 and 18 °C, has been purified without bacterial contaminants (Fig. 5b and 5c). A single band (∼34 kDa) in the eluted fractions was observed on SDS-PAGE, indicating the high purity of HAND2 recombinant fusion protein [Fig. 5b (*top*), 5c (*top*), 5d and 5e]. This is crucial because purified proteins having bacterial proteins are likely to induce undesired effects on the target human cells when applied to elucidate their biological function [49]. However, loss of HAND2 fusion protein was detected in the immunoblotting analysis (Fig. 5b and 5c) in the flow-through/unbound fraction. This could probably be due to the overloading of the soluble fraction on the purification column or the low volume of resin used for purification. Further, this fusion protein identity was established by immunoblotting with the HAND2 antibody [Fig. 5b and 5c (*bottom*)]. Interestingly, the purified HTN-HAND2 protein was eluted efficiently only from the second elution fraction (Fig. 5d and 5e), signifying that the interactions between the protein molecules and Ni-NTA were stronger and requires more elutions with elution buffer to weaken these interactions before eluting. The elution profile of one-step affinity-purified HAND2 fusion proteins induced at 37 °C and 18 °C is shown in Fig. 5f and Fig. 5g, respectively. The purification data are summarized in Table 2, and the final yield of the purified HAND2 fusion protein was around 0.87 and 0.60 mg/g of wet cells when induced at 37 and 18 °C, respectively. Thus, the one-step purification of recombinant HAND2 fusion protein under native conditions from soluble fractions was demonstrated, irrespective of the induction temperatures.

**Fig. 5.**
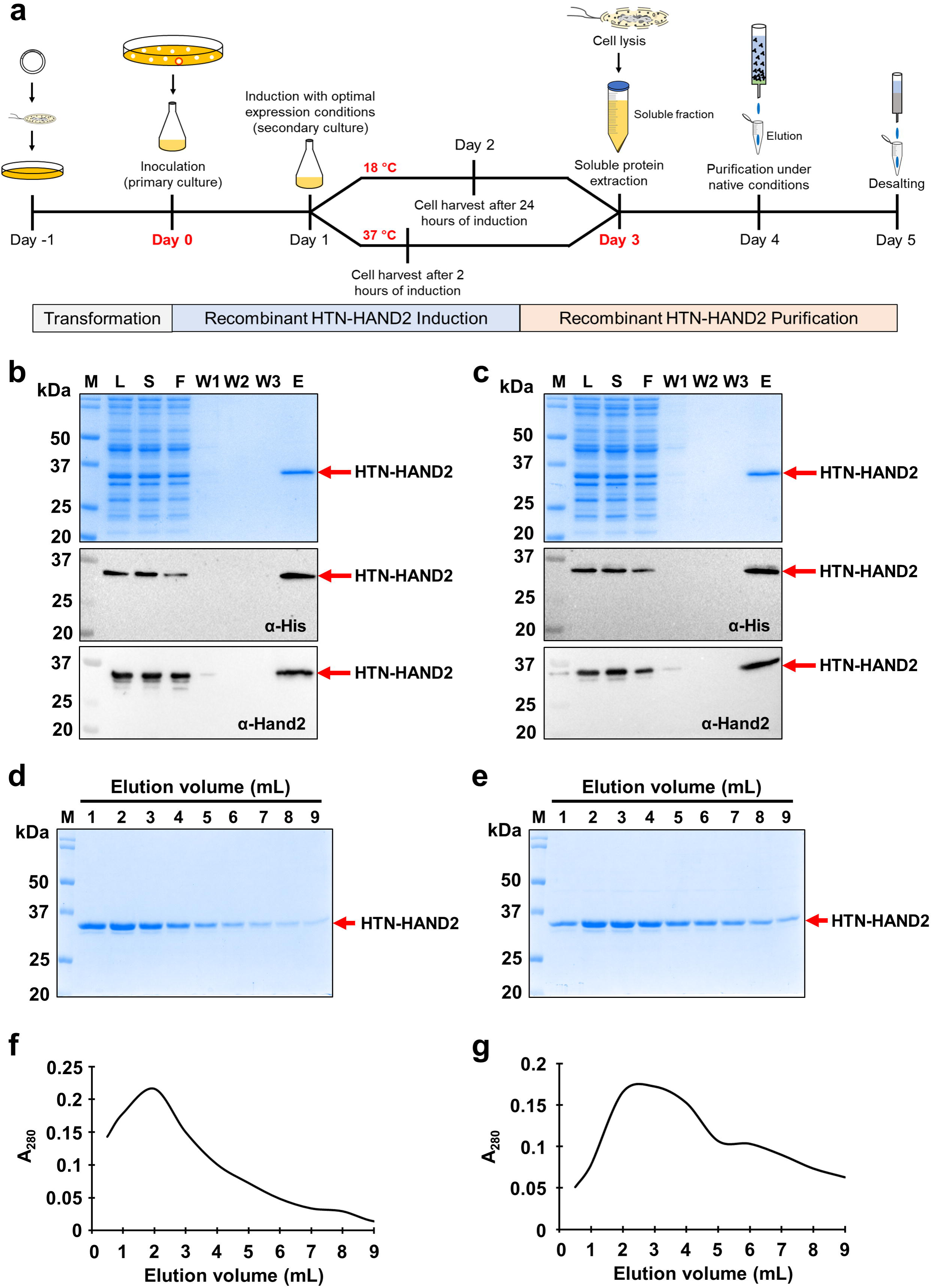
One-step purification of recombinant human HAND2 protein, induced at two different temperatures. **a** Timeline and pictorial representation of the overall experimental strategy. **b** and **c** The purification of HTN-HAND2 protein at two different induction temperatures 37 and 18 °C, respectively. The samples collected during different stages of purification were analyzed using SDS-PAGE (*top*) and immunoblotting using an α-his (*middle*) and α-HAND2 (*bottom*) antibody. 20 µg/lane for L fraction, equal volumes for S-W3 fractions corresponding to L fraction, and 40 µL of E fraction were used for the analysis. **d** and **e** Analysis of elution fractions of the purified HAND2 fusion protein induced at 37 and 18 °C, respectively. The expressed HTN-HAND2 protein, induced at two different temperatures, was purified, and the eluted proteins were analyzed using SDS-PAGE gel with equal loading volume. **f** and **g** Elution profile of one-step homogeneous purification of the HAND2 fusion protein induced at 37 and 18 °C, respectively. M: protein marker, L: lysate, S: soluble supernatant, F: flow-through, W1: wash buffer 1, W2: wash buffer 2, W3: wash buffer 3, E: elution.

**Table 2.** Purification summary of recombinant human HTN-HAND2 protein

| Induction<br>temperature<br>(°C) | Steps | Total protein | Target protein | Yield | Purity |
| --- | --- | --- | --- | --- | --- |
|  |  | (mg)<sup>b</sup> | (mg)<sup>c</sup> | (%) | (%)<sup>d</sup> |
| 37 | Crude lysate <sup>a</sup> | 58.89 | 10.40 | 100 | 17.67 |
|  | Cleared lysate (soluble) | 55.30 | 9.88 | 95 | 17.87 |
|  | IMAC (pooled) | 0.88 | 0.87 | 4.13 | 99.01 |
| 18 | Crude lysate <sup>a</sup> | 54.65 | 9.13 | 100 | 16.70 |
|  | Cleared lysate (soluble) | 50.76 | 7.65 | 83.79 | 15.07 |
|  | IMAC (pooled) | 0.62 | 0.60 | 5.37 | 97.92 |

<sup>a</sup> From 1 g wet weight of induced *E. coli* BL21(DE3) cell pellet (from 0.25-0.35 L of culture).
<sup>b</sup> Protein concentration determined by Bradford assay using BSA as a standard protein.
<sup>c</sup> Determined from total protein concentration and purity.
<sup>d</sup> Purity determined from SDS-PAGE analysis using ImageJ software

### MALDI-TOF and secondary structure analysis of human HTN-HAND2 protein

The monoisotopic molecular weight of the recombinant HAND2 fusion protein 30687.805 Da was calculated from the amino acid sequence using Peptide Mass Calculator online server (https://www.peptidesynthetics.co.uk/tools/). The molecular weight was accurately determined using MALDI-TOF mass spectrometer and found to be 30.6 kDa (Fig. 6a). The difference in the observed and calculated molecular weight remains well within the calibration error of the instrument. We next sought to study the secondary structure content of the purified recombinant HAND2 protein, as the crystal structure of this human protein has not been reported yet. Commonly, the CD spectroscopic technique is used to study the folding conformation/characteristics of desired proteins whose secondary structure is unknown [50, 51], like human HAND2. Principally, different secondary structures absorb different amounts of right and left circularly polarized light. Therefore, the spectrum obtained from the CD technique is characteristic in shape and magnitude to the specific type of secondary structure of the protein, namely α-helices, β-sheets, turns, and random coils [50, 51]. Typically, in a CD spectrum, α-helices show two negative peaks at 222 and 208 nm and a single positive peak at 193 nm [50]. Correspondingly, β-sheets have a negative peak at 218 nm and a positive peak at 195 nm, whereas random coils have a negative peak at 195 nm and a positive peak at 210 nm [50]. The overall spectra are obtained depending on the various amounts of secondary structure content present in the given protein.

**Fig. 6.**
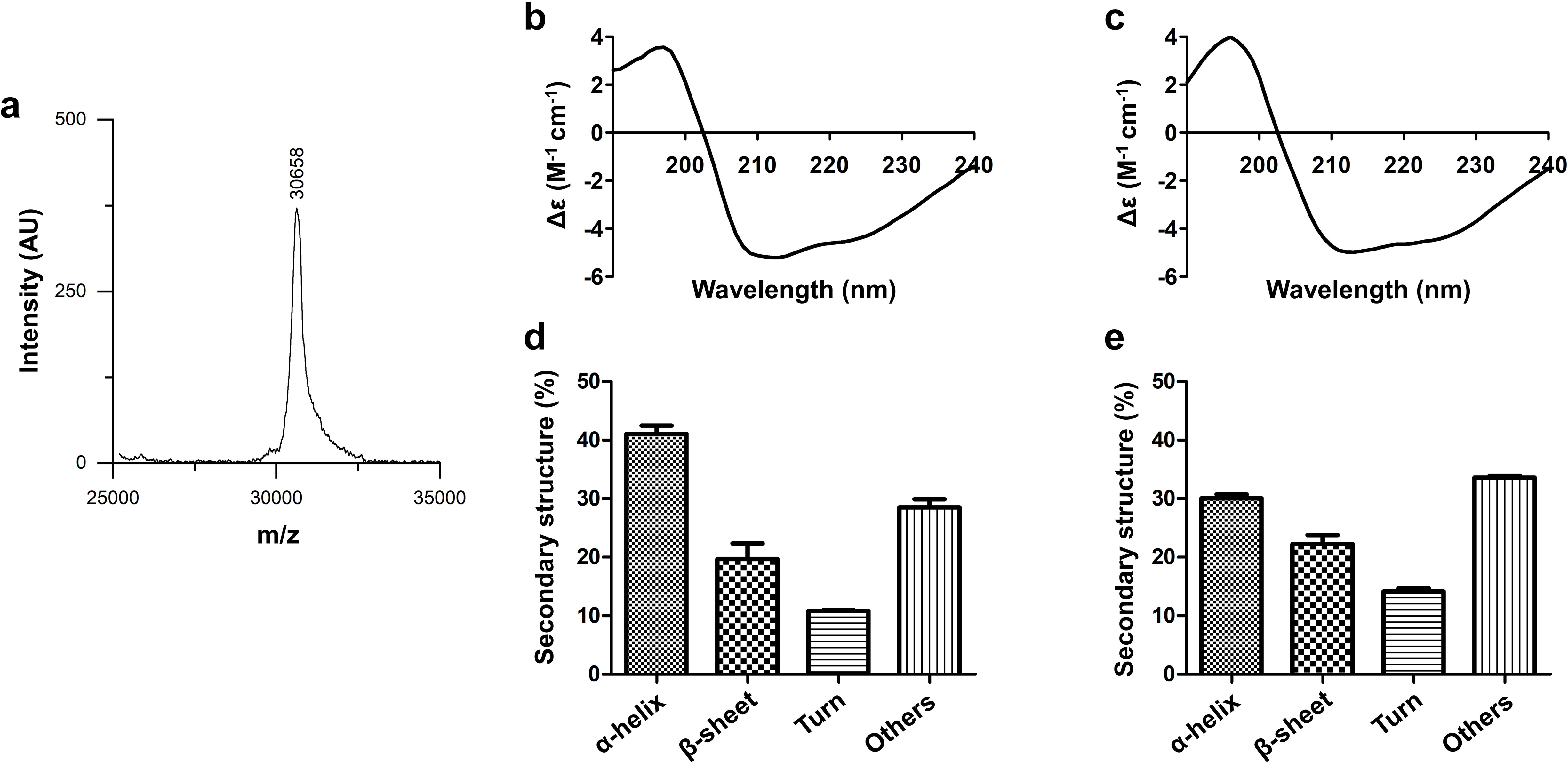
MALDI-TOF and Circular dichroism spectroscopy analysis of the purified recombinant HAND2 fusion protein. **a** The MALDI-TOF-MS analysis of the purified HAND2 protein. The purified HAND2 protein was analyzed using MALDI-TOF, and the result is depicted using a graph with mass per charge (m/z) ratio in the x-axis and intensity (AU) on the y-axis. **b-e** The secondary structure content of this purified HAND2 fusion protein was determined using far-UV CD spectroscopy. The obtained far-UV CD spectra were then evaluated using *in silico* BeStSel online tool. **b** and **c** The CD spectrum of the purified HAND2 fusion proteins induced at 37 and 18 °C, respectively, were plotted with Delta Epsilon (M^-1^ cm^-1^; Y-axis) against wavelength (nM; X-axis). **d** and **e** The bar graphs represent the quantified secondary structure of the purified HTN-HAND2 proteins induced at 37 and 18 °C, respectively.

From the CD spectra of purified recombinant human HAND2 protein, induced at two different temperatures, 37 °C and 18 °C (Fig. 6b and 6c), it is evident that HAND2 protein has maintained its secondary structure. Moreover, the level of similarity between both spectra indicates that the secondary structure is consistent irrespective of the induction temperature. Subsequently, from the CD spectra, different secondary structures were quantified using BeStSel online tool. The secondary structure content of HAND2 protein, induced at two different temperatures, 37 °C and 18 °C (Fig. 6d and 6e), reveals that the purified protein comprises majorly of α-helices and other structures (mostly random coils). From the results (Fig. 6d and 6e), it is apparent that 41% of α-helices and 28% of other structures are observed when the induction temperature was 37 °C, whereas 30% of α-helices and 34% of other structures are observed when the induction temperature was 18 °C. Furthermore, there is a significant contribution of β-sheets (20% (induced at 37 °C) and 22% (induced at 18 °C)) and turns (11% (induced at 37 °C) and 14% (induced at 18 °C)) to the secondary structure of HAND2 protein. Although the spectra appeared similar, variability in the secondary structure content was observed upon the quantification using BeStSel analysis. This variability is likely due to the induction and purification of HTN-HAND2 at different temperatures, which might have resulted in differences in their folding. Also, the possible minor differences in the protein concentrations (37 °C v/s 18 °C) may also be the reason for this variability. However, due to the non-availability of the 3D structure of human HAND2 protein, the spectra and the secondary structure content determined in this study may be slightly different from the native protein or the secondary structure content of the native protein may be similar to either 37 °C or 18 °C. Further comparison can be made once the crystal structure is available. In addition, the presence of fusion tags may also alter the structure of the HAND2 protein, and thereby the secondary structure content estimated in this study may be slightly different from the native protein. In general, our results have established that this purified HTN-HAND2 protein has maintained its secondary structure post purification.

### Effect and transduction ability of recombinant form of the purified HAND2 fusion protein in human cells

Further, we investigated the transduction ability of this purified fusion protein in human cells. BJ fibroblasts and HeLa cells were exposed either with 200 nM of recombinant HTN-HAND2 protein, induced at two different temperatures, 37 and 18 °C, or glycerol buffer as a negative control (vehicle control) for 4-6 h and then analyzed using fluorescence microscopy (Fig. 7a and 7b). Two different cell lines, BJ and HeLa, were chosen for protein transduction analysis because they lack endogenous expression of HAND2 protein. Also, BJ human fibroblasts were selected since they are widely used for cellular reprogramming studies [13, 14], and HeLa cells are commonly used for transduction studies. As shown in Fig. 7a and 7b, the transduction of HAND2 fusion protein across the sub-cellular and sub-nuclear region of both cell lines was observed. The microscopy results further confirmed the absence of endogenous expression of HAND2 in both BJ fibroblasts and HeLa cells as shown in the vehicle control panel, and this also signifies that the glycerol buffer does not trigger the expression of HAND2 or did not lead to any false positive signal during analysis (Fig. 7a and 7b). Thus, this purified recombinant fusion protein (irrespective of the induction temperature in which they were expressed) has cell penetration (mediated by TAT) and nuclear translocation (mediated by NLS) ability, indicating the functionality of these tags post-purification.

**Fig. 7.**
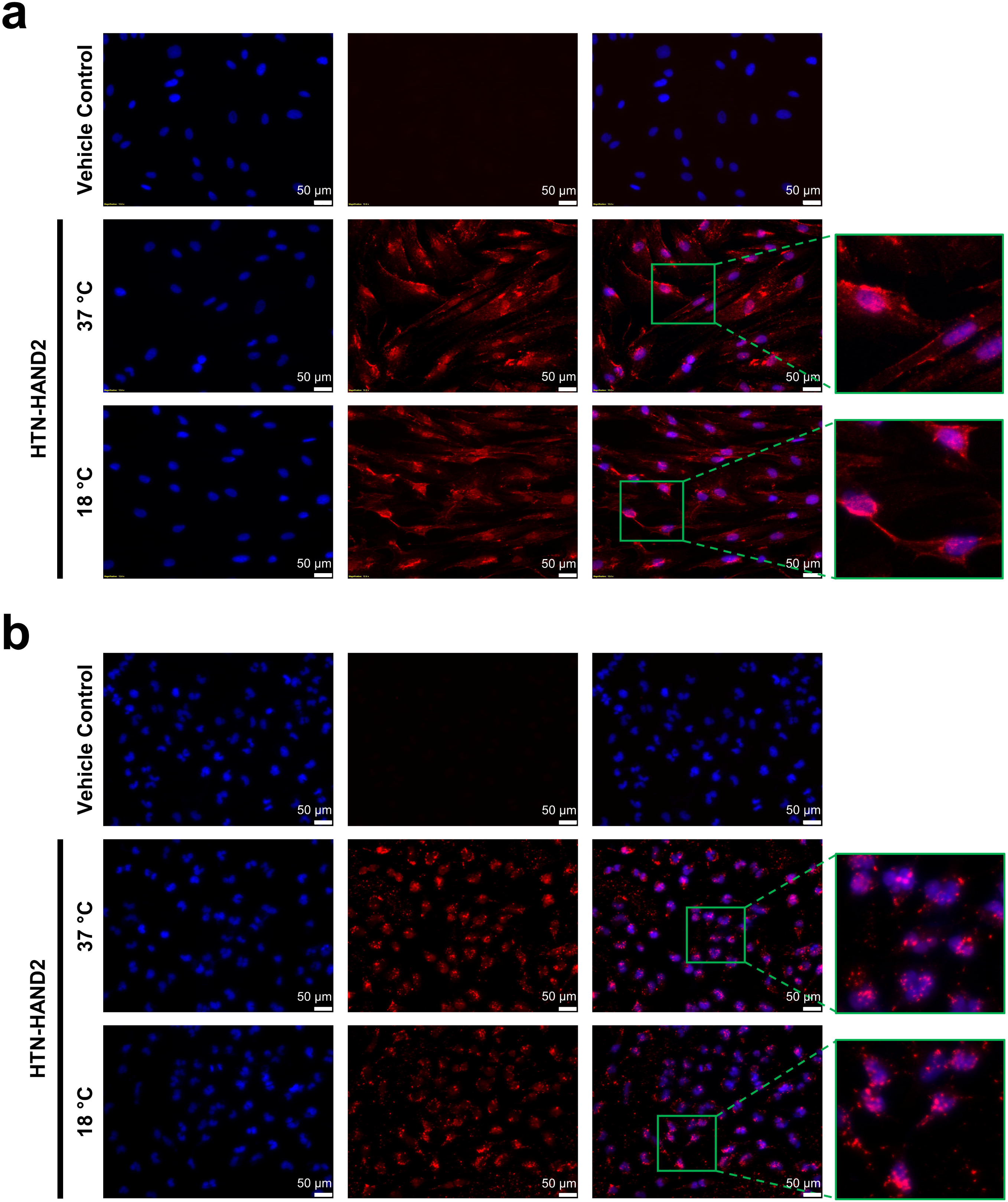
Subcellular and subnuclear delivery of purified recombinant HTN-HAND2 protein into human cells. HFF (**a**) and HeLa (**b**) cells were exposed to 200 nM of purified recombinant HTN-HAND2 protein for 4-6 hours at standard cell culture conditions. Subsequently, cells were fixed, permeabilized, and blocked, and then stained with an anti-HAND2 antibody. Transduced cells were detected with an Alexa Fluor® 594 conjugated anti-rabbit secondary antibody. Nuclei were stained with DAPI, and images were taken at 20x magnification. Scale bar: 50 μm (n=3)

## Discussion

Here we demonstrate the purification of a transducible version of the recombinant human HAND2 fusion protein, a crucial cardiac reprogramming factor to derive functional myocytes from the non-myocytes. Several studies have reported that the inclusion of this transcription factor HAND2 in GATA4, MEF2C, and TBX5 cocktail improved the cardiac reprogramming efficiency [25–27]. However, all these studies used the genetic form of this reprogramming factor and delivered it through viral vectors to derive induced cardiomyocytes, thereby limiting its therapeutic applications. On this account, generating a transducible HAND2 protein is crucial to obviate the potential risks associated with using viruses and DNA transfection in cardiac reprogramming.

First and foremost, we codon-optimized the *HAND2* protein-coding sequence to express it in the heterologous system efficiently. Several studies on multiple genes have demonstrated the enhanced heterologous expression of human genes in *E. coli* after codon optimization [43, 45, 52], eliminating the roadblocks associated with the protein expression. In recombinant protein production, eliminating the codon bias appears to be one of the critical factors contributing to its success [53, 54] either by co-expressing tRNAs encoding genes or by optimizing the coding sequence [43, 45, 52]). The latter choice has a more positive impact on the heterologous expression of human genes in the bacterial expression systems, unlike the tRNA-enhanced host systems [43, 45, 52]). Apart from eliminating codon bias, codon optimization also removes destabilizing RNA elements, alters GC content to appropriate levels, avoids RNA secondary structures, removes direct repeats, eliminates cryptic splice-sites, avoids intragenic poly(A)-sites, and so forth [43, 45, 52]. Thus, we have codon-optimized the *HAND2* gene for its expression in *E. coli*, and consistent with our previous studies [29-31, 33, 34]; here we also found an increased CAI value of codon-optimized *HAND2* sequence than its non-optimized sequence. Results further established that the *HAND2* sequence is absent in any rare codons after codon optimization.

The expression host used in this study is *E. coli*. It has a high transformation efficiency and is economical for recombinant protein production in large quantities, fast growth, and well-known genetics [52, 55]. In addition, this organism is commonly used for human recombinant protein production that does not require posttranslational modifications for functionality [56–61]. Notably, phosphorylation of HAND2 by Akt leads to the formation of the heterodimer, thereby inhibiting its transcriptional activity [62]. Therefore, to circumvent this modification, particularly *E. coli* strain BL21(DE3) (lacks *lon* and *OmpT* proteases) was used for the stable expression of human HAND2 fusion proteins. Also, this strain is enhanced with T7 RNA polymerase, compatible with the pET28a(+) expression vector, and ideal for the overexpression and stability of the desired protein [63].

In the present study, we report for the first-time screening and identification of the ideal expression parameters for the soluble expression of recombinant HAND2 fusion proteins in *E. coli*. Consistent with this, we and others have previously demonstrated the effect of expression parameters on the expression, solubility, stability, and secondary structure conformation of recombinant proteins [29-31, 33, 34, 37-43]. Moreover, we also observed the effect of tagging fusion tags at either terminal in the expression and production of the quality recombinant HAND2 fusion protein, similar to our previous observations with ETS2 and MESP1 recombinant proteins [30, 31]. To the best of our knowledge, this is the first study to establish a one-step purification to obtain a highly pure recombinant human HAND2 fusion protein under native (from soluble cell fraction) conditions that has its secondary structure retained. Notably, it can transduce into the mammalian cells and translocate to its nucleus without a transduction reagent. Similar strategies have been reported earlier by numerous studies for recombinant transcription factors purified from *E. coli*, namely, OCT4, NANOG, SOX2, PDX1, ETS2, GATA4, and MESP1 [13, 29–31, 33–35, 64, 65]. Our strategy of coupling this transcription factor with the protein transduction domain and NLS facilitated its delivery into the mammalian cell target site. As a transcription factor, it has its major role inside the nucleus; therefore, its nuclear entry is one of the most critical aspects of its functionality. Similar fusion strategies were employed in previous studies, including us for the efficient subcellular and subnuclear delivery of reprogramming factors such as OCT4, NANOG, SOX2, PDX1, and GATA4 in the form of recombinant proteins in mammalian cells [29, 33, 35, 64–67]. These studies have used TAT and NLS as fusion tags to efficiently deliver protein of interest into the cell and its nucleus, respectively, to exert their biological function in mammalian cells [13, 35, 64, 65].

Notably, this transducible version of recombinant HAND2 fusion protein does not require any additional protein transduction reagent for its effective delivery to the target site in mammalian cells [13, 68]. Previous studies have employed cell-permeant recombinant transcription factors in the cell reprogramming process and established that their functionality is comparable to their genetic forms [66, 67]. Most importantly, in all these studies, the presence of the fusion tags (TAT and NLS) did not hinder their functionality. Hence, this purified HAND2 recombinant protein can be utilized to generate transgene-free cells as well as for other biological applications; thus, eliminating the limitations that accrue due to the plasmid or viral-based approaches [69, 70, 71]. In summary, we have successfully conducted codon optimization, molecular cloning and expression, and demonstrated one-step homogeneous purification of the full-length human HAND2 reprogramming factor from *E. coli* in the form of recombinant protein. The established approach is simple, cost-effective, and highly reproducible. Further, we have identified the optimal expression conditions and showed the impact of the fusion tags coupled at either terminal in the recombinant HAND2 fusion protein expression. Besides, our highly pure recombinant version of HAND2 protein had retained its folding conformation and confirmed its ability to transduce the cells as well as translocate to the nucleus. Amongst the plethora of possible biological applications of the recombinant HAND2 fusion protein, it can also be used as a substitute against its viral and genetic form in direct cardiac reprogramming process to investigate its mechanistic role.

## Compliance with ethical standards

### Ethical approval

This article does not contain any studies with human participants or animals performed by any of the authors.

### Author contribution

**KKH** was responsible for conception and design, collection and/or assembly of data, data analysis and interpretation, manuscript writing, and final approval of the manuscript; **PKS** was responsible for collection and/or assembly of data, data analysis and interpretation, manuscript writing, and final editing and approval of the manuscript; **AM**, **NJ** and **NJD** were responsible for collection and/or assembly of data, data analysis and interpretation, and **RPT** was responsible for conception and design, collection and/or assembly of data, data analysis and interpretation, manuscript writing, final approval of the manuscript and financial support. All the authors gave consent for publication.

## Acknowledgments

We thank all the members of the Laboratory for Stem Cell Engineering and Regenerative Medicine (SCERM) for their critical reading and excellent support. We thank Dr. Kusum K Singh Lab, IIT Guwahati for providing HeLa cells, and Dr. Shirisha Nagotu, IIT Guwahati, for the use of fluorescence microscopy facility for this study. This work was supported by North Eastern Region – Biotechnology Programme Management Cell (NERBPMC), Department of Biotechnology, Government of India (BT/PR16655/NER/95/132/2015), and also by IIT Guwahati Institutional Top-Up on Start-Up Grant (RPT). KKH, PKS, AM, and NJ acknowledges the Ministry of Education (MoE), Government of India, for providing fellowship.

## Supplementary figure captions

**Figure S1** Codon optimization and cloning workflow.

**Figure S2** Analysis of *HAND2* gene sequences before and after codon optimization using GRCA online tool. The codons with values ≤ 30 (a dotted line indicates threshold) are more likely to diminish the expression of HAND2 in *E. coli*.

**Figure S3** Analysis of HAND2 gene sequences before and after codon optimization using GCUA tool. Codons with a relative adaptiveness value ≤ 30% (*magenta*) are likely to diminish the expression of HAND2 in *E. coli*.

