## Supplementary figures and images for "Generation of transducible version of a recombinant human HAND2 transcription factor from *Escherichia coli*"

### Figure S1

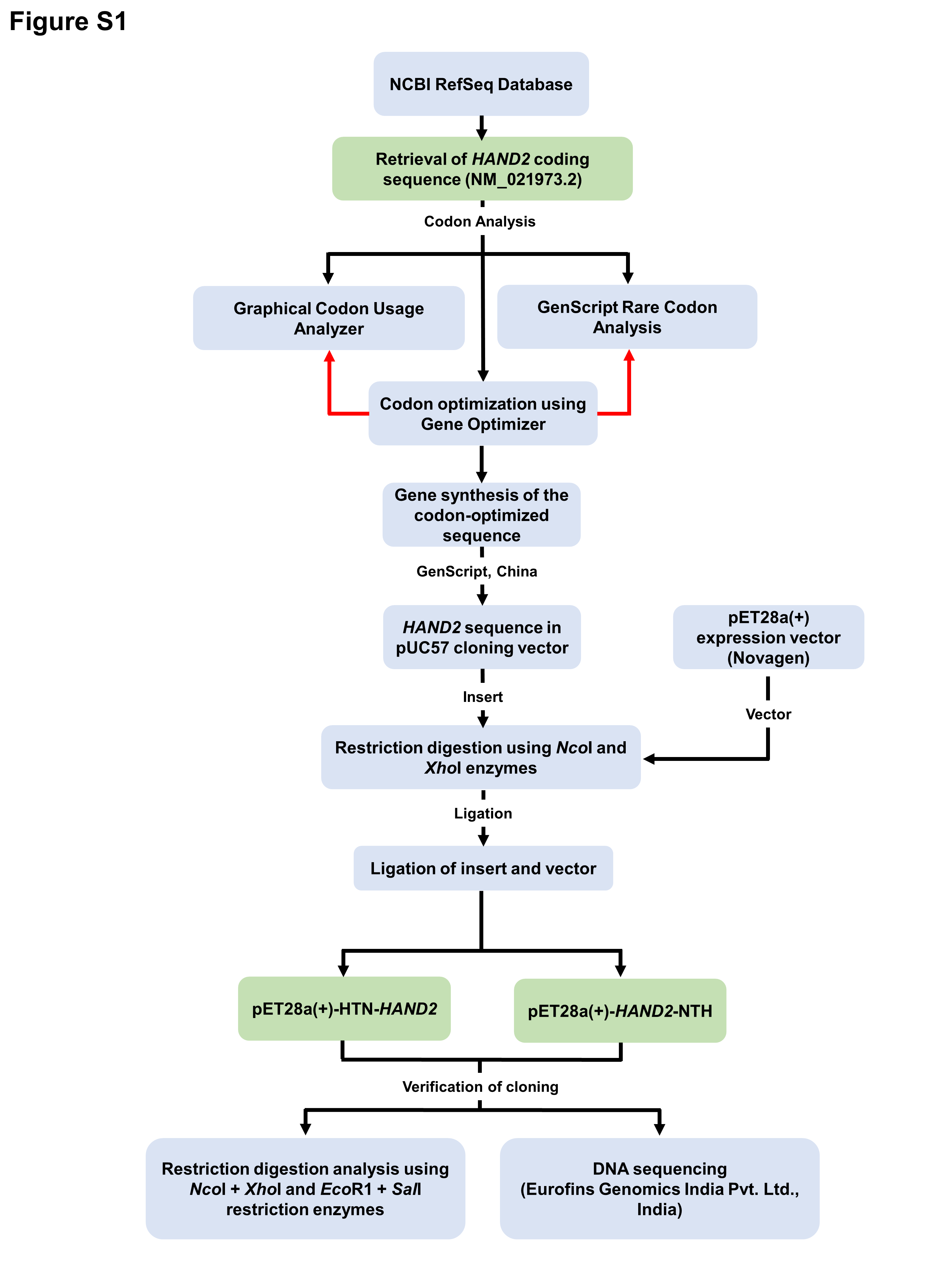

### Figure S2

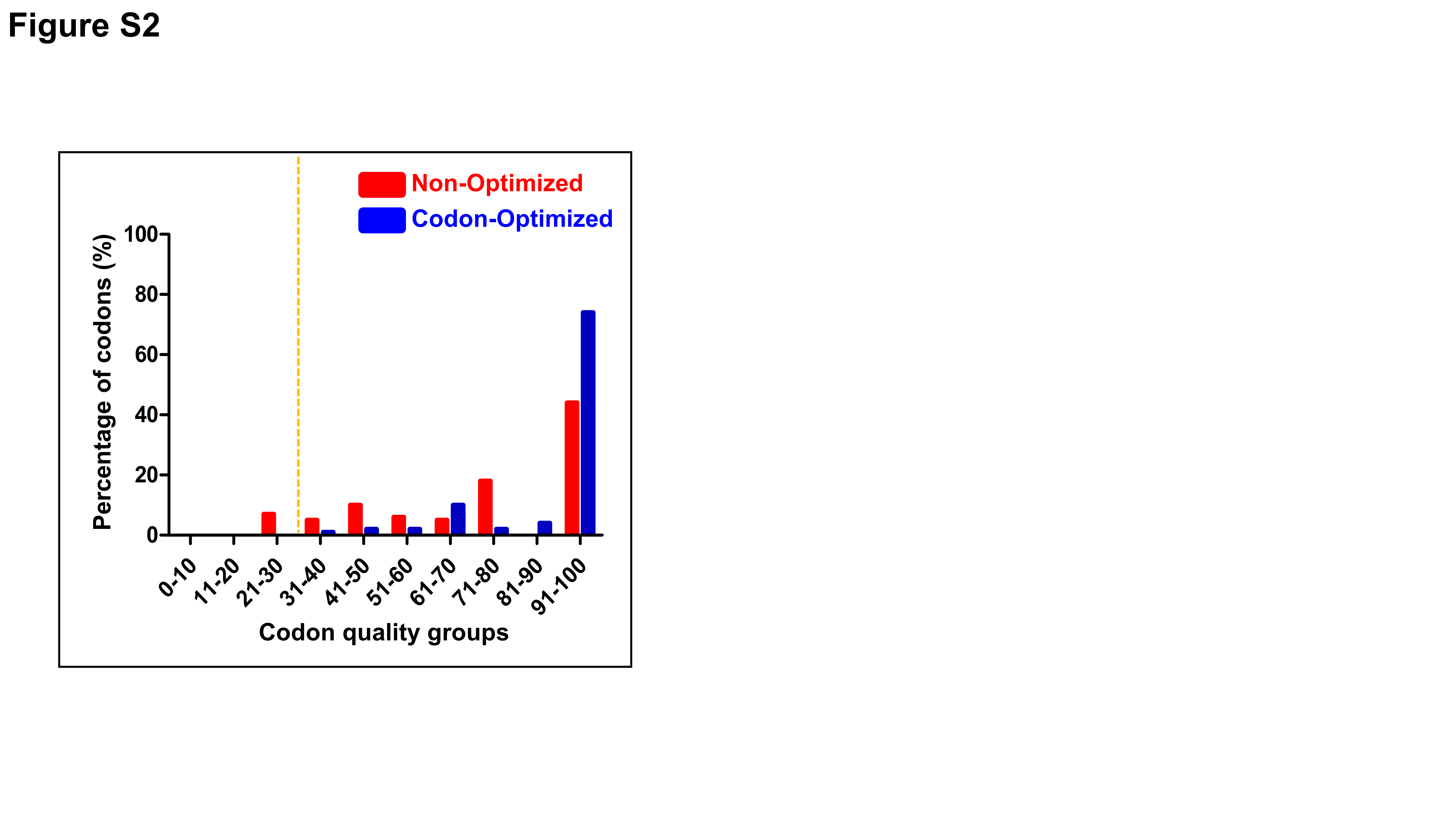

### Figure S3

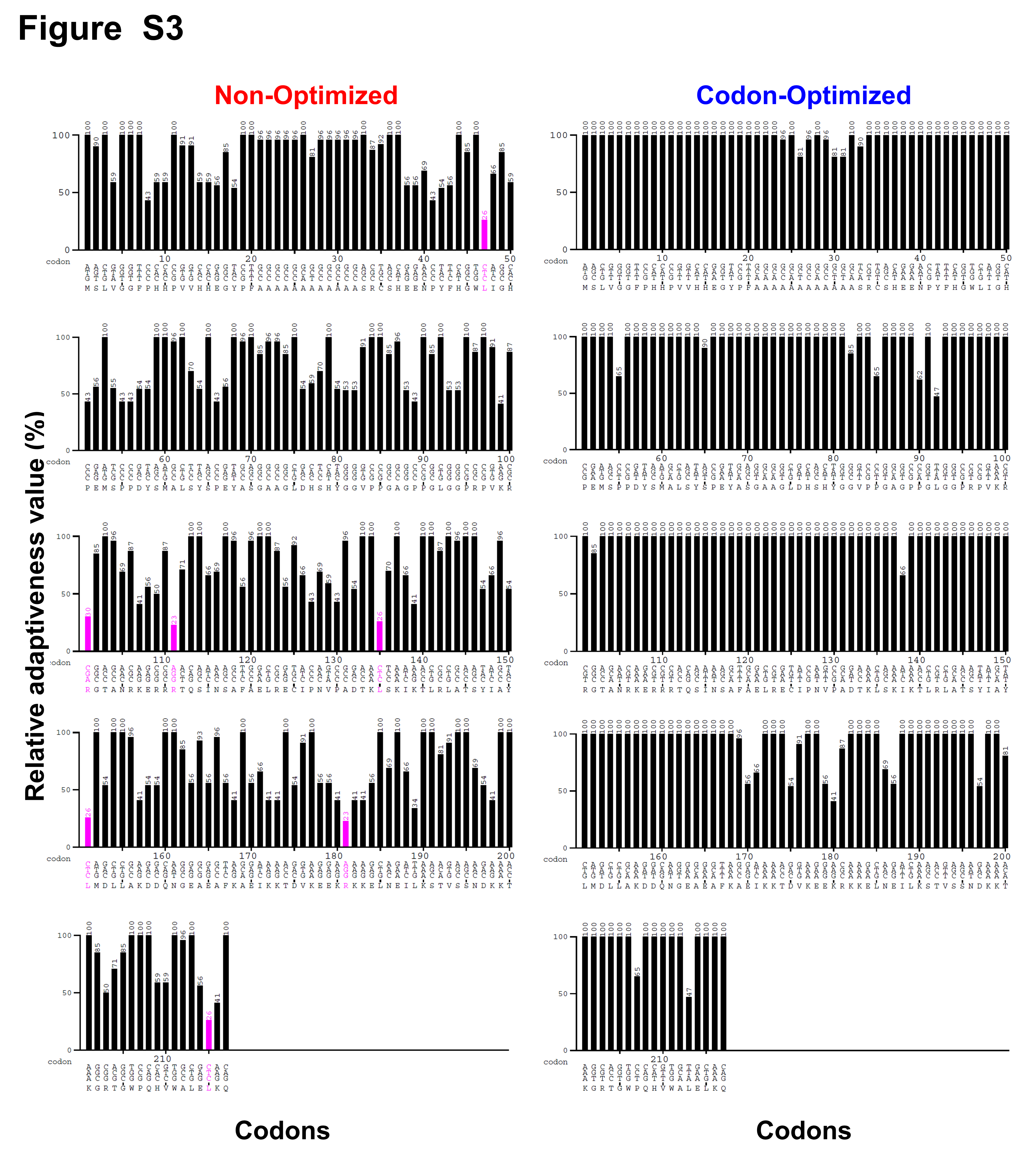
