## Supplementary material for "Generation of transducible version of a recombinant human HAND2 transcription factor from *Escherichia coli*": Table S1

**Table S1** List of *in silico* online tools used for codon optimization and evaluation

| Online tools | Weblink | Reference(s) |
|---|---|---|
| **Codon optimization** | | |
| ThermoFisher Scientific GeneOptimizer | https://www.thermofisher.com/in/en/home/life-science/cloning/gene-synthesis/geneart-gene-synthesis/geneoptimizer.html | |
| **Evaluation of optimized codons** | | |
| Graphical Codon Usage Analyser 2.0 | http://gcua.schoedl.de/ | |
| Genscript Rare Codon Analysis Tool | https://www.genscript.com/tools/rare-codon-analysis | |
