## Supplementary material for "Generation of transducible version of a recombinant human HAND2 transcription factor from *Escherichia coli*": Table S2

**Table S2** Primers used for verification of gene inserts by DNA sequencing

| **Primers** | **Sequence (5’ – 3’)** |
|---|---|
| Forward Primer | TAATACGACTCACTATAGGG |
| Reverse Primer | GCTAGTTATTGCTCAGCGG |
