## Supplementary material for "Generation of transducible version of a recombinant human HAND2 transcription factor from *Escherichia coli*": Table S3

**Table S3** List of expression parameters and their values screened in expression analysis

| Parameters | Values to be screened |
|---|---|
| Inducer concentration (mM) | 0.05, 0.1, 0.25, and 0.5 |
| Pre-induction cell density ($OD_{600}$) | 0.5, 1.0, and 1.5 |
| Post-induction temperature (°C) | 37 and 18 |
| Post-induction incubation time (h) | For 37 °C: 2, 4, and 8 |
| | For 18 °C: 12, 24, and 48 |
