## Supplementary material for "Generation of transducible version of a recombinant human HAND2 transcription factor from *Escherichia coli*": Table S4

**Table S4** Summary of purification buffers used and their composition

| Ingredients | Lysis buffer | Wash buffer | | | Elution buffer | Glycerol buffer |
|---|---|---|---|---|---|---|
| | | W1 | W2 | W3 | | |
| Phosphate buffer (mM) | 20 | 20 | 20 | 20 | 20 | 20 |
| NaCl (mM) | 150 | 150 | 150 | 150 | 150 | - |
| Imidazole (mM) | 20 | 50 | 100 | 150 | 300 | - |
| Glycerol (%) | 20 | - | - | - | 20 | 20 |
