## Supplementary material for "Generation of transducible version of a recombinant human HAND2 transcription factor from *Escherichia coli*": Table S5

**Table S5** List of antibodies used in this study

| Antibodies | Dilutions | | Company | Identifier |
|---|---|---|---|---|
| | Western blotting | Immunostaining | | |
| Anti-His | 1:5000 | N/A | BioBharati | Cat # BB-AB0010 |
| Anti-Hand2 | 1:2000 | 1:40 | Invitrogen | Cat # PA5-35186 |
| Anti-Rabbit IgG, HRP-conjugated | 1:5000 | N/A | Invitrogen | Cat # 31460 |
| Alexa Fluor 594 goat anti-rabbit IgG | N/A | 1:1000 | Invitrogen | Cat # A11037 |
