## Supplementary material for "Generation of transducible version of a recombinant human HAND2 transcription factor from *Escherichia coli*": Table S6

**Table S6** *HAND2* fusion gene constructs and expected size after its restriction digestion

| Vector/inserts/constructs | Restriction Enzyme(s) | Expected Size (bp) |
|---|---|---|
| pET28a(+) Empty Vector | - | 5369 |
| *HAND2* | - | 654 |
| HTN-*HAND2* / *HAND2*-NTH | - | 839 |
| pET28a(+)-HTN-*HAND2* / pET28(+)-*HAND2*-NTH | - | 6064 |
| pET28a(+)-HTN-*HAND2* / pET28(+)-*HAND2*-NTH | *Nco*I and *Xho*I | 833 and 5231 |
| pET28a(+)-HTN-*HAND2* / pET28(+)-*HAND2*-NTH | *Eco*RI and *Sal*I | 657 and 5407 |
